# Acoustic detection of a rarely vocalising invasive mammal from sparse data

**DOI:** 10.64898/2026.06.19.733324

**Authors:** Anthony Gibbons, Andrew Parnell, Ian Donohue, Masako Ogasawara, Samuel R. P.-J. Ross

## Abstract

1. Monitoring and limiting the spread of invasive species on islands requires efficient detection and population estimation methods. However, elusive species can be difficult to monitor using traditional methods, making autonomous approaches such as camera trapping and acoustic monitoring increasingly valuable.
2. On the island of Okinawa, Japan, the small Indian mongoose ( *Urva auropunctata*) threatens many native species since its introduction in 1910. Listed among the world’s worst invasive species, effective monitoring of U. *auropunctata* in Okinawa is critical. The Okinawa Environmental Observation Network (OKEON) uses camera traps to detect *U. auropunctata*, but success depends on precise placement. Though OKEON also includes a high-resolution acoustic monitoring programme, no audio classification model currently exists for *U. auropunctata*. Developing such a model could improve substantially our capacity to detect and manage the species.
3. Using sparse *U. auropunctata* vocalisations collected from camera trap videos, we built a lightweight Convolutional Neural Network distilled from a more complex model for classifying contact calls and alarm calls of *U. auropunctata*. Our distilled model performed similarly to the full model at detecting vocalisations from training data, but was considerably faster.
4. We applied the distilled classifier to ∼486 hrs of audio collected over eight years from southern Okinawa, where we successfully detected *U. auropunctata* a handful of times in each year of recording. In spite of strong model performance on test data, our model did not transfer well to unseen data, perhaps owing to the rarity of *U. auropunctata* calls and consequent small training dataset size, limiting its utility for ecological monitoring.
5. *Practical implication*. The use of sparse audio data from camera trap videos to train an acoustic classifier had limited utility to detect the rarely vocalising *U. auropunctata* from passive acoustic monitoring data. We provide several recommendations for enhancing classifier performance to provide robust actionable insights into the distribution and spread of *U. auropunctata*, and aid targeted conservation efforts for Okinawa’s threatened biodiversity.

## 1 Introduction

Invasive species significantly threaten biodiversity globally, disrupting ecosystem functions and services through predation, competition, and habitat alteration (Roy et al. 2024). Often introduced through human activity, invasive species can establish rapidly, particularly on islands where native species have small populations and limited prior exposure to novel interactions with invasive species (Reaser et al. 2007). For example, the brown tree snake *(Boiga irregularis*) has driven several bird species to extinction on Guam in the Mariana islands (Wiles et al. 2003), and invasive rats drive declines in tropical island populations by preying on native bird eggs and chicks (Harper and Bunbury 2015). At the same time, islands are often hotspots of endemism, yet island ecosystems are among the worst affected by human impacts including land use change and invasive species (Kier et al. 2009; Reaser et al. 2007). Accordingly, endemic species on islands face a mounting threat from invasive species (see e.g. Borges et al. 2006). The global economic cost of invasive species damage exceeds US$100 billion annually (Diagne et al. 2021), underscoring the urgency of early detection and effective management strategies, especially in vulnerable island ecosystems.

Recent advances in scalable biodiversity monitoring provide promise for detecting invasive species and mitigating their devastating ecological impacts (Gasc et al. 2017; Hofstadter et al. 2022). Passive acoustic monitoring (PAM) has emerged as a powerful tool for assessing biodiversity, offering a non-invasive, scalable, and cost-effective approach to studying sound-producing animals. By deploying autonomous microphones, PAM records species vocalisations to infer species presence, diversity, and ecological dynamics for a range of purposes (Ross, O’Connell, et al. 2023; Stiff, Wakefield, and Rands 2026; Soh et al. 2024). For example, PAM has been used to monitor bat activity (Mac Aodha et al. 2018), document bird migration (Van Doren et al. 2023), track individual bird species population trends (Wood et al. 2020; Borker et al. 2014), reveal freshwater ecosystem dynamics (Gottesman et al. 2020) and, notably, to identify and monitor invasive species (Rountree and Juanes 2017). However, the sheer volume of audio data produced by PAM studies poses its own challenge, particularly when searching for infrequent events or rarely vocalising species (Gibb et al. 2018). Recent advances in machine learning, particularly deep learning-based image classification models, have gained popularity in bioacoustics for species classification of animal vocalisations. These classifiers vary in complexity, ranging from simple presence /absence detection (Mac Aodha et al. 2018) to those that identify a few (Manriquez et al. 2024), dozens (Zhong et al. 2020; Gibbons, King, et al. 2024), or even hundreds (Kahl et al. 2021; Ghani et al. 2023) of distinct species. Such pre-trained models can be applied to different situations but often require labelled training data for fine-tuning to specific use cases.

The small Indian mongoose ( *Urva auropunctata*,フィ リマングース in Japanese) was introduced to Okinawa Island in 1910 by Professor Shozaburo Watase to control venomous pit vipers *(Protobothrops flavoviridis*, ハブ)and agricultural pests. Watase believed strongly in human intervention and had witnessed a mongoose preying on a cobra in Sri Lanka, informing his decision to introduce *U. auropunctata* as an “ecological experiment” (Kaneko 2021). Without contemporary knowledge of ecological balance or consideration of long-term consequences, his introduction of *U. auropunctata* cast a long shadow over Okinawa. Today, the small Indian mongoose is one of the world’s worst invasive species (Lowe et al. 2000), and Okinawa is a hotspot for cross-taxon species invasions (Wayne Dawson et al.2017). In Okinawa, the mongoose began spreading northward, reaching Yambaru forest by the 1990s (Yamada and Sugimura 2004). Yambaru forest is a biodiversity hotspot, home to a rich fauna and flora including species endemic to Okinawa island (McWhirter et al. 1996; Ito, Miyagi, and Ota 2000; Miyamoto, Tamanaha, and Watari 2021). *U. auropunctata* has had devastating ecological consequences for various small mammals, reptiles, amphibians, and insects (Environmental Risk Research Center, National Institute for Environmental Studies, Japan 2025), but particularly for Okinawa’s rich avifauna, exacerbating population declines of the critically endangered Okinawa rail(*Gallirallus okinawae*,ヤンバルクイナ),Okinawa woodpecker *(Dendrocopos noguchii*,丿グチゲラ),and Okinawa robin *(Larvivora namiyei*,アカヒゲ )(Yagihashi et al. 2021), with indications that these species are still declining (see Figure S1).

Efforts to mitigate the mongoose’s impact began with a removal project in Yambaru forest (2000-2004) and a formal control plan under Japan’s Invasive Alien Species Act of 2004 (https://www.env.go.jp/en/nature/as.html). Control measures include live traps and bird-excluding kill traps, with mongoose control fences erected in 2005–2006 to isolate a 280 km^2^ area in Yambaru for targeted eradication. The Okinawa Prefectural Government and Japan’s Ministry of the Environment currently aim for complete eradication of the mongoose inside the fenced area by 2027. *Urva auropunctata* capture rates are declining in Yambaru (https://kyushu.env.go.jp/okinawa/press_00109.html) and the government recently declared complete extirpation from Amami-Oshima Island, North of Okinawa (https://www.env.go.jp/en/press/press_03205.html). Despite these efforts, the mongoose population in Okinawa is still significant, particularly outside of the fence where intervention has been minimal. As such, scalable methods for detection and monitoring are needed to measure the success of future conservation intervention.

Here, we use an eight-year acoustic monitoring dataset from outside of the mongoose control fence in Okinawa to investigate the efficacy of using PAM to monitor the rarely vocalising invasive mongoose, *Urva auropunctata*. We built a mongoose vocalisation classifier using sparse audio data extracted from camera trap videos, and applied this classifier to audio collected at a single site of the Okinawa Environmental Observation Network (OKEON) (Ross, Friedman, Dudley, Yoshimura, et al. 2017; Ross, Friedman, Dudley, Yoshida, et al. 2023). We intend our acoustic classification model to complement traditional *U. auropunctata* population surveys in Okinawa, providing additional evidence for the presence and distribution of *U. auropunctata* in future.

## 2 Materials and Methods

### Environmental Monitoring Programme

This study uses data from the OKEON (Okinawa Environmental Observation Network) Churamori Project (OKEON 美ら森プロジェクト in Japanese; https://www.oist.jp/core-facilities/marine-terrestrial). OKEON aims to monitor terrestrial biodiversity through time at 24 field sites distributed across a range of Okinawa’s terrestrial habitats (Figure 1a). Each field site deploys modified malaise traps to capture insects, environmental sensors for weather data, and Song Meter SM4 acoustic recording units (Wildlife Acoustics Inc., Concord, MA, USA) and Hyperfire 2 Covert camera traps (Reconyx Inc., Holmen, WI, USA) to monitor birds, mammals, and other taxa (Stiff, Wakefield, and Rands 2026). Camera traps are installed at a height of ∼1.0 m and are movement-triggered. Acoustic recorders are fixed at ∼1.3 m at each site, facing North, and record with two omnidirectional microphones at default gain (+16 dB) on a duty cycle of 1-min recording, 29-min standby. Recording starts every hour and half-hour, and audio data are saved to SD cards in stereo .WAV format with a sampling rate of 48 kHz, then converted to lossless .flac format. Camera traps and acoustic sensors have been installed since 2017, with previous work in Okinawa demonstrating the efficacy of such remote monitoring techniques for detecting endemic birds, bats, and rats (Kobayashi et al. 2022; Ross, Friedman, Dudley, Yoshimura, et al. 2017; Dinets et al. 2020; Yamada, Kawauchi, et al. 2010), and broadly understanding biodiversity and disturbance (Ross, Friedman, Yoshimura, et al. 2021; Ross, Friedman, Dudley, Yoshida, et al. 2023).

**Figure 1.**
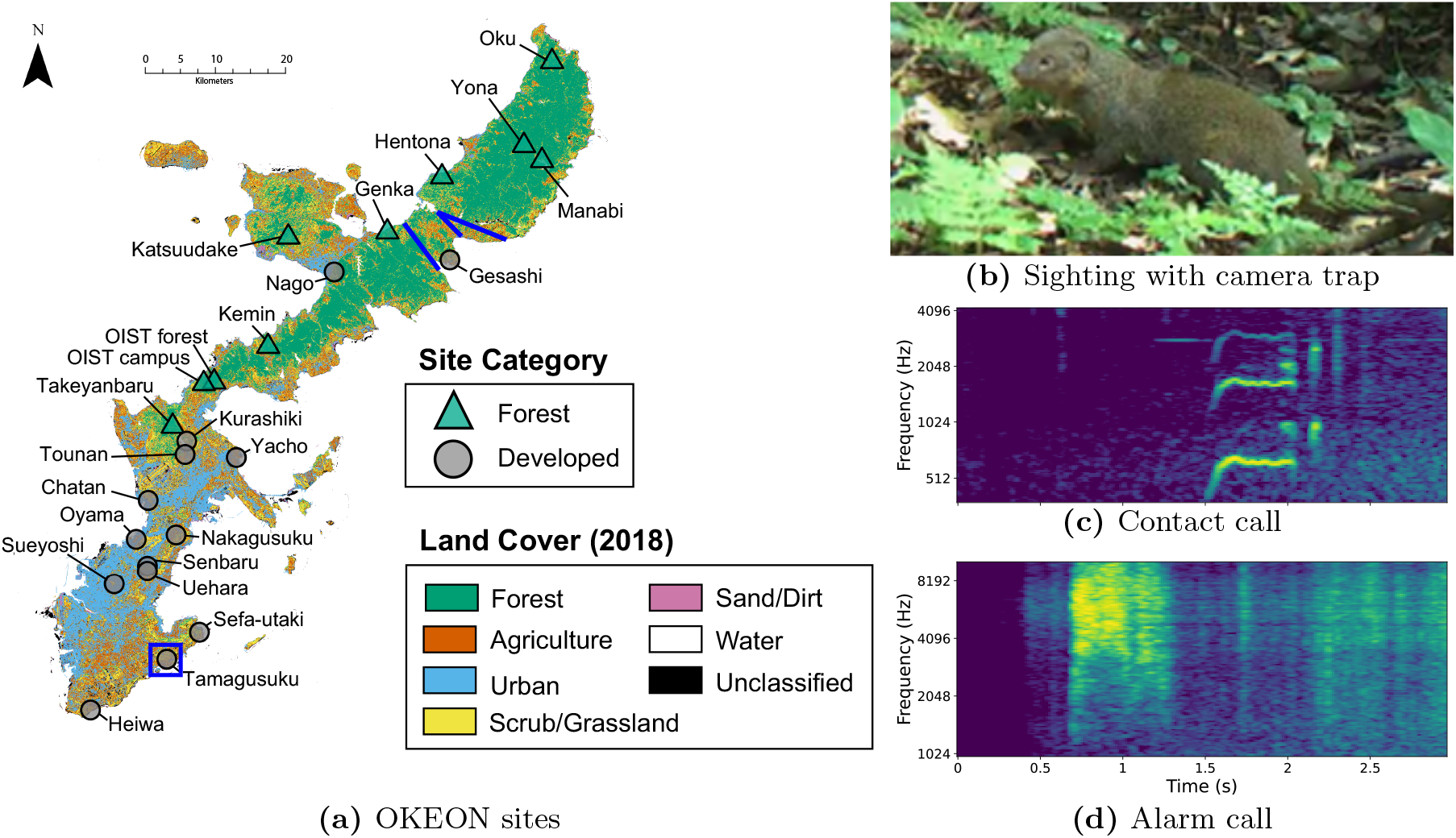
OKEON, camera trap still and example extracted sounds. (a) Map of OKEON monitoring sites across Okinawa, including land use classification from (Ross, Friedman, Dudley, Yoshimura, et al. 2017). Tamagusuku, our focal field site, is highlighted by the blue box, and mongoose control fences are located around the blue lines. (b) Still *U. auropunctata* image from movement-triggered camera trap video. (c,d) Spectrograms of audio from *U. auropunctata* contact (c) and alarm calls (d). Mongoose alarm calls tend to occupy higher frequencies than contact calls; note the differing vertical axes on panels (c) and (d).

Here, we combine camera trap video (Figure 1b) with PAM audio to build a mongoose vocalisation classifier. Detection of animals using movement-triggered camera traps depends on precise camera placement, whereas PAM sensors cover a much broader detection area with omnidirectional microphones, providing higher relative detection efficiency and probability (Enari et al. 2019; Crunchant et al. 2020). However, *U. auropunctata* rarely vocalises—1.28% of 2,500 manually screened *U. auropunctata* video samples included vocalisations (see below)—producing only occasional contact calls, mainly between pups and adults (Figure 1c) and alarm calls when fighting conspecifics or capturing prey (Figure 1d). Accordingly, initial detection and verification are easier using camera trap data than audio, while audio recordings complement targeted field surveys with wider spatial coverage (Darras et al. 2018). To initially screen for mongoose vocalisations to use as training data, we used 4,099 movement-triggered camera trap videos from 2020 and 2022 at 20 OKEON field sites. The videos totalled 9.7 hours (5.7 GB), with individual files ranging from 6-10 seconds (mean 8.5 s). Almost all sample files contained *U. auropunctata*.

### Data Preparation and Annotation

Approximately 2,500 of 4,099 camera trap videos (Figure 2a) were screened manually for *U. auropunctata* vocalisations, yielding 32 examples, which were supplemented with 7 examples from 2025. We also selected an equal number of ‘empty’ clips from the many confirmed not to contain mongoose audio. Twenty more examples were found via semi-supervised learning (see Supplementary Information for detailed description), for a total of 52 vocalisations from 2022 (Figure 2b). Once verified, we extracted audio from camera trap videos using MoviePy (Zulko et al. 2025). Unless otherwise stated, all analyses used audio data. Another 11 example audio files (mostly alarm calls) were web-scraped from YouTube, Facebook and Xeno-Canto (see Table S1). In total, our search and screening process produced 70 examples of mongoose vocalisations. Most mongoose detections were during the hours 07:00-18:00 and vocalisation density was highest during late Summer (∼July-September; Figure 2b). *Urva auropunctata* was detected most often at the Tamagusuku OKEON site in the Southeast of Okinawa (Figure 2c).

**Figure 2.**
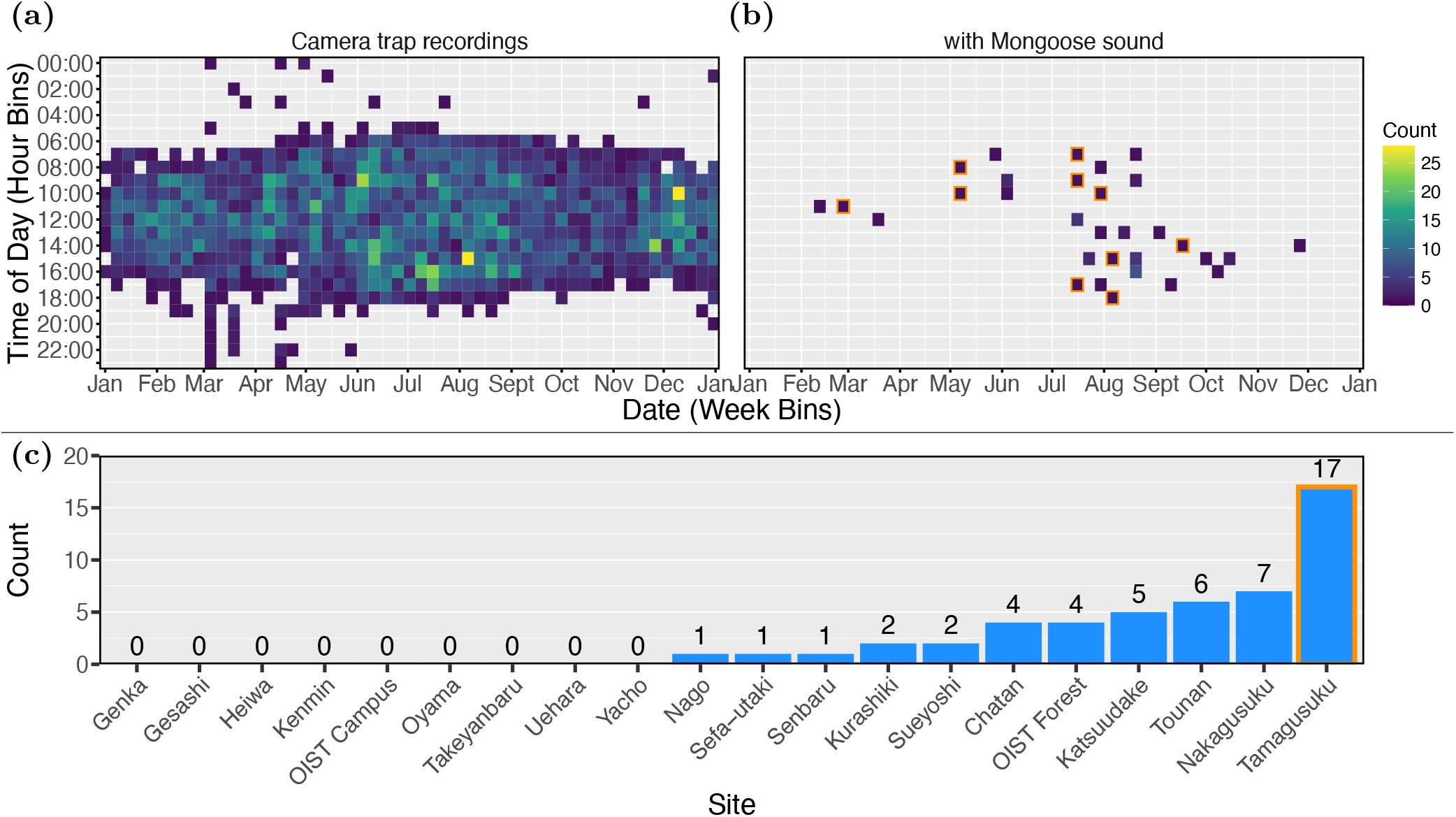
Counts of camera trap and acoustic detections by day and hour across all sites. (a,b) The y-axis shows the hour of day (0:00-23:00) and the x-axis is broken into week bins ranging from 1 January to 31 December 2022, inclusive. Brighter colours represent higher validated *U. auropunctata* counts in video (a) and audio (b) from camera traps. Few detections are between 19:00 and 05:00, or in the months November-May. (c) *Urva auropunctata* vocalisation counts from camera trap audio per OKEON site across the sound clips. Tamagusku, outlined in orange in both plots, represents the site with the majority of mongoose vocalisations from camera traps, and thus was selected as our acoustic monitoring case study because of its higher likelihood of containing *Urva auropunctata* vocalisations in scheduled recordings.

We labelled the 70 mongoose audio clips using NEAL (Gibbons, Donohue, et al. 2023), finding 134 distinct vocalisations. We enriched the dataset by copying mongoose clips and combining them with clips from the inference dataset via mixup, resulting in 244 mongoose examples. We took many more examples without mongoose vocalisations from the remaining clips and a sample from the inference dataset to better balance the mongoose-to-non-mongoose ratio, making the dataset more representative of the target area. The final classification dataset had 2,005 empty and 244 mongoose examples. The training and test sets were split randomly (80% training, 20% test), stratified by label. Examples were then reassigned to the training set if they were web-scraped, contained only a single mongoose vocalisation, or had a filename already appearing in training. To better reflect inference conditions,1,039 additional empty examples from OKEON acoustic data were added to training and 54 to test. To carry out fine-tuning without overfitting to the test set, the test set was split randomly in two. The final training-validation-test ratio was 0.92:0.04:0.04. The resulting training set contains 3,644 empty and 196 mongoose samples. The validation and test sets each include 197 empty (non-mongoose) and 28 mongoose samples. To classify mongoose vocalisations, we used both VGGish, which looks directly at the audio, and ResNet, which looks at the 256x256 spectrogram images. See Supplementary Information for more detail.

### Species Classification Models

Convolutional Neural Networks (CNNs) have become a highly effective tool for image classification over the last decade (Krizhevsky, Sutskever, and Hinton 2012), including for bioacoustic species classification from spectrograms (Stowell 2022). CNNs perform well, learning task-specific features directly, and are robust to noise and variability in field recordings (Lostanlen et al. 2019). Moreover, researchers can leverage transfer learning from pre-trained models to reduce the training data required to reach adequate performance with their CNNs (Zhao et al. 2024). We first compared the performance of two CNNs of different complexities (Table 1). The larger, teacher model uses transfer learning of two pre-trained CNNs: VGGish and ResNet. VGGish uses raw audio (Hershey et al. 2017) and ResNet uses spectrogram images (He et al. 2015), providing complementary but computationally complex information for audio classification. To address the trade-off between generalisation and computational intensity, we also built a smaller, student classifier to be trained using knowledge distillation, where the larger model from the first iteration serves as the teacher (Hinton, Vinyals, and Dean 2015). This student CNN takes only spectrograms as input and does not directly use the pre-trained classifiers, but retains much of the teacher’s predictive power while reducing the number of parameters and inference time as needed for scaling to large audio datasets (Table 1). For further details of teacher and student models, see Supplementary Information.

**Table 1.** Comparison of Models. The teacher model takes in both a 3-channel spectrogram and VGGish embedding as inputs, while the student model uses a single-channel spectrogram. The ratio column shows how much greater each metric is for the teacher model relative to the student model. Given its extra preprocessing steps of the ResNet and VGGish runs, the teacher model has over 10 times the parameters, 100 times the multiply-add operations to carry out and is over 10 times larger in storage than the student model. If a student model approaches the teacher’s performance, the student model would be advantageous for scaling to large datasets.

| Metric | Teacher Model | Student Model | Ratio |
| --- | --- | --- | --- |
| **Input Shape** | [(3, 256, 256), (384,)] | (1, 256, 256) | - |
| **# Parameters** | 12 060 650 | 1 090 050 | 11.06 |
| **# Multiply-Adds** | $2.38 \times 10^9$ | $2.23 \times 10^7$ | 106.73 |
| **Total Size (MB)** | 92.77 | 8.29 | 11.19 |

We evaluated classifier performance using several metrics: accuracy (proportion of correct predictions), precision (true positives divided by all predicted positives), recall (true positives divided by all actual positives), F1 score (harmonic mean of precision and recall), and the Area Under the Receiver Operating Characteristic Curve (AUC) score, which plots the true positive rate against the false positive rate at different detection thresholds. Collectively, these metrics allow us a broad understanding of model performance (Rainio, Teuho, and Klen 2024). Both models used the same training hyperparameters: Adam optimiser, learning rate of 0.001,a batch size of 16, and 50 epochs.

In machine learning, a loss function quantifies the discrepancy between a model’s predictions and the true outcomes, guiding the optimisation process during training. For species classification, categorical cross entropy measures the difference between the model’s probabilistic output and the true class labels, while Kullback-Leibler (KL) divergence (Kullback and Leibler 1951) compares two probability distributions (e.g. between two models) and a cosine embedding loss (PyTorch Core Team 2024) measures the similarity between two embeddings (outputs from earlier layers of the model). The teacher model in this study used cross entropy loss while the student model, employing knowledge distillation, used cross entropy as the target loss and the sum of KL divergence (with temperature *T* = 3) and cosine embedding loss (with margin m = 0) as the distillation loss, with a weighting *a* = 0.7 on distillation loss and 1—*a* on target loss. Total loss is the sum of the two. For further details of the loss functions used, see Supplementary Information. We trained a copy of the student model, which we call the vanilla student model, using just categorical cross entropy loss and not by distilling the teacher model. We did this to evaluate performance when training from scratch, without taking advantage of learnings from the pre-trained models.

The hardware environment for our model comparison was a local machine with 16 GB of RAM and an Intel(R) Core(TM) i7-1165G7 CPU @ 2.80 GHz, with 4 cores and 8 threads. The operating system was Windows 11 Pro 24H2. The programming environment included Visual Studio Code (version 1.99.3) (Microsoft Corporation 2025) for prototyping, training, and post-processing, using Python (version 3.9.21) (Python Software Foundation 2024) and PyTorch (version 2.6.0) (PyTorch Team 2025) within this environment.

### Acoustic Monitoring Case Study

We applied the chosen model to a long-term audio dataset collected as part of the Okinawa Environmental Observation Network’s passive acoustic monitoring programme (Ross, Friedman, Dudley, Yoshimura, et al. 2017). We chose one OKEON field site, Tamagusuku, as our case study for out-of-sample model inference, owing to the relative number of mongoose vocalisations in camera trap audio from this site (Figure 2). We chose a single site because the data size, over 150 GB per site across the 24 sites on the island, was too large to process for all sites in the OKEON dataset. Tamagsuku forest is a small disturbed forest surrounded by urban land use (Figure 1a). The forest patch has alkaline soil, supporting typical Okinawan flora such as *Ficus microcarpa* (ガン ユマ丿レ),*F. superba* (アコウ),and *F. virgata* (ノハマイ ヌ ビワ)which germinate and grow on rocks and large trees. The tree canopy is primarily composed of *Cinnamomum yabunikkei* (ヤブニッ ケイ)and *Celtis boninensis Koidz* (クワ丿ハエ丿キ),with other climbing plants such *Flagellaria indica* (トウツレモドキ) and *Ipomoea indica* (丿 アサガオ),and palm trees including *Arenga engleri* (クロ ッ グ).Despite the small forest patch size and anthropogenic noise from the surrounding city, Tamagusuku also has a rich soniferous fauna, including resident songbirds such as the Japanese tit *(Parus cinereus*,シン ユ ウカラ),and migratory species like the Japanese paradise flycatcher (*Terpsiphone atrocaudata*,サ ンコウチョウ).The site is also home to endemic frogs *(e*.*g., Rhacophorus viridis*,オキナワアオガ エル),newts *(e*.*g., Cynops ensicauda popei*,シュリ ケンイモリ),and lizards *(e*.*g., Goniurosaurus kuroiwae*,クロイワ トカゲモドキ).Several species of small bats roost in nearby limestone caves, but do not produce sound in the focal frequency range of our study.

We used audio recordings corresponding to peak mongoose vocalisation periods from preliminary data (Figure 2); we targeted 07:00-19:00 each day between 15 May and 14 October from 2017 to 2024. Taking each 1-minute audio file per half hour (25 files/day) each Summer (153 days/year) for eight years resulted in 29,159 min (∼486 hrs) of continuous audio totalling 155 GB compressed to .flac format. We ran the distilled model over this data, and manually verified *Urva auropunctata* detections labelled with > 0.8 confidence.

## 3 Results

### Evaluating model performance

The large teacher model achieved 94.6% accuracy on the test data, using the default threshold of 0.5, with mongoose-specific metrics of 0.61 precision, 0.92 recall, 0.73 F1-score and AUC of 0.96. The vanilla student model was trained similarly, using just the spectrograms as inputs and species labels as outputs, and achieved 91.9% accuracy with 0.70, 0.61, 0.65 and 0.92 for precision, recall, F1-score and AUC, respectively. The distilled student model, trained both on the target labels and outputs from the teacher model achieved best validation loss, achieved 92.6% accuracy, with 0.63 precision, 0.78 recall, 0.80 F1-score and 0.99 AUC. See figure 3 for detail.

**Figure 3.**
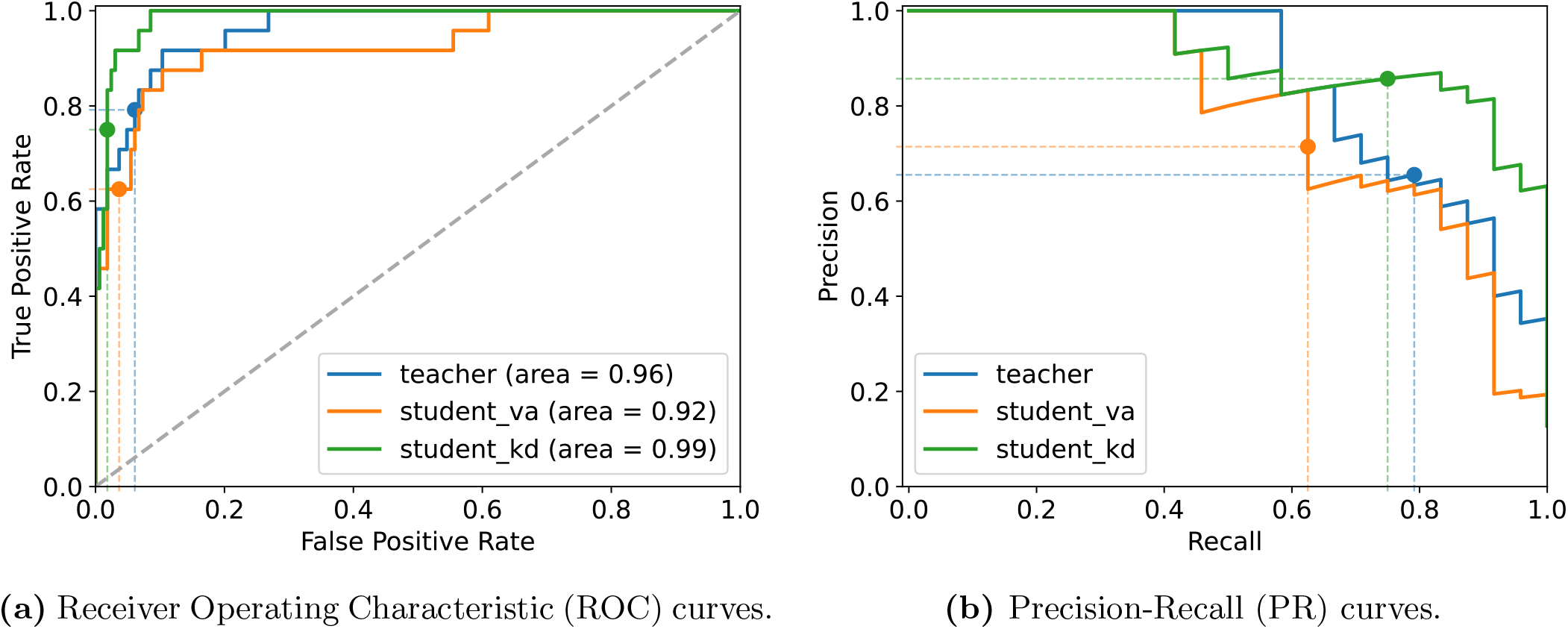
Performance comparison of teacher and student models. (a) Receiver Operating Characteristic (ROC) curve for each model, showing how well they distinguish between correct and incorrect detections. (b) Precision-Recall (PR) curves, illustrating the trade-off between the two metrics. Curves were created by adjusting the detection threshold from 0 to 1 (default = 0.5, shown at points and coloured dashed lines) and analysing output probabilities for *Urva auropunctata* in the test set. For (a), we calculated true positive and false positive rates, and for (b), we measured precision and recall at each threshold. The grey dashed line on the diagonal in (a) is a random guess, where false positive and false negative rates are expected to be the same at any threshold. **student_va**, the vanilla student model, is the smaller model architecture trained solely on the labelled data, while **student_kd** is the same architecture trained via knowledge distillation using outputs from the **teacher** model.

The distilled model shows significantly better metrics than the vanilla student model, and even surpasses the teacher model at most thresholds. Coupled with its computational efficiency 10-fold higher than the teacher model, we opted to apply the distilled model to our full dataset. We first trained the distilled model again on all the data (training, validation, and test) to improve its generalisation. This retraining used knowledge distillation to transfer some of the learnings from the large teacher model to a more computationally efficient architecture.

### Application to monitoring data

We applied our trained model to the large acoustic dataset to classify mongoose vocalisations. We ran the model on each year of the dataset, processing audio recordings to identify mongoose and non-mongoose (empty) vocalisations. Model outputs included considerably more non-mongoose classifications than predicted mongoose vocalisations (Figure 4).

**Figure 4.**
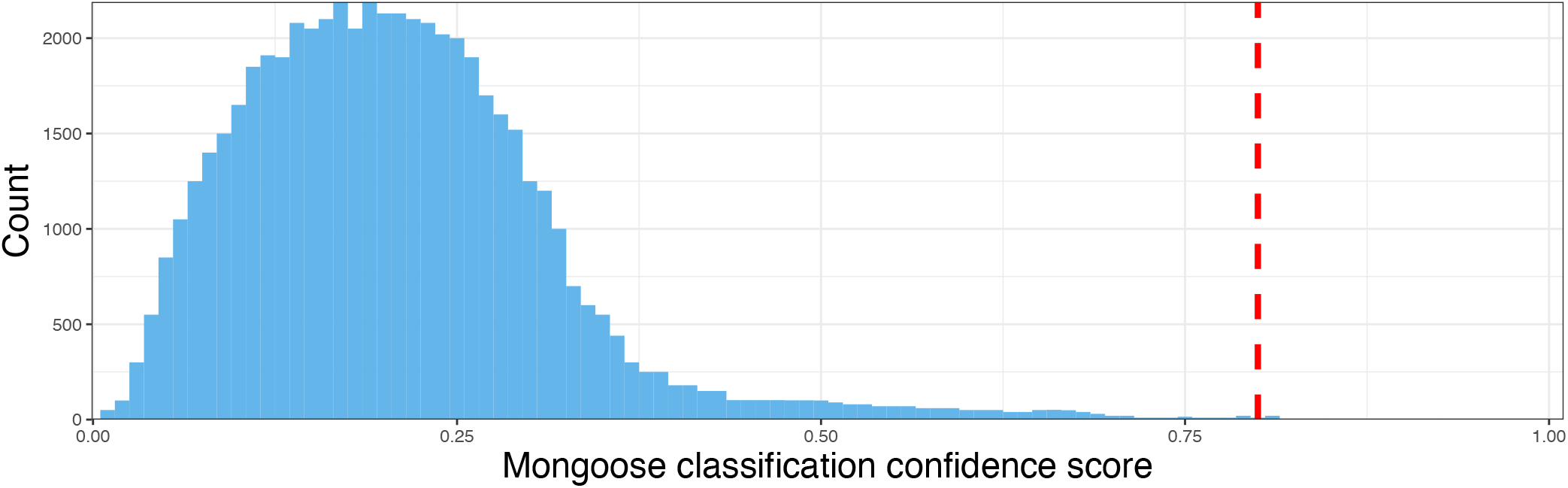
Histogram of mongoose classification confidence from 2022 data. The model predictions (confidence scores) are noticeably skewed left, with mostly low-confidence predictions (0.1-0.4 confidence), below the default 0.5 cutoff. The distribution has a sharp decline after 0.4, and a long tail towards 1.0. This is consistent with mongoose being a rarely vocalising species not often present in audio recordings. Of the 62,039 predictions from 2022, only 54 had confidence over 0.8 (red dashed line).

We manually verified model predictions above a 0.8 confidence threshold and identified a total of 5 mongoose vocalisations across the entire dataset (Table 2). Despite the limited number of verified samples among the predictions (0.66% precision across all years), these serve as a supplemental examples for expanding the training set for semi-supervised learning. See Figure S4 for a detailed visualisation of the model’s confident predictions through time.

**Table 2.**
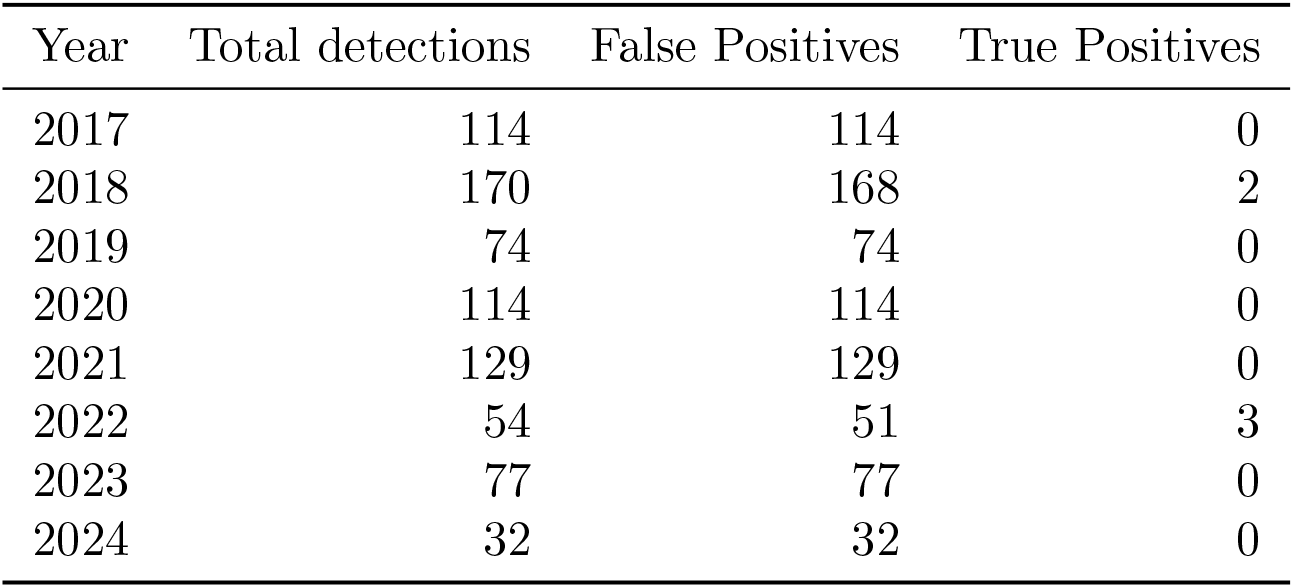
High-confidence mongoose detections by year. Number of model predictions with >0.8 confidence score per year and verified true positives versus false positives.

| Year | Total detections | False Positives | True Positives |
| --- | --- | --- | --- |
| 2017 | 114 | 114 | 0 |
| 2018 | 170 | 168 | 2 |
| 2019 | 74 | 74 | 0 |
| 2020 | 114 | 114 | 0 |
| 2021 | 129 | 129 | 0 |
| 2022 | 54 | 51 | 3 |
| 2023 | 77 | 77 | 0 |
| 2024 | 32 | 32 | 0 |

To analyse the model’s behaviour on the spectrograms in the inference data, we examined saliency maps—or class activation maps—to better understand which time-frequency regions and patterns most strongly influence model predictions (Zhang et al.2021). These maps are created using gradients of the model’s output score with respect to the input spectrogram, and are computed via backpropagation to identify influential pixels in the image. They highlight regions of the spectrogram image that contribute most to classification decision-making, thereby allowing us to interpret whether the model is using meaningful acoustic features or is overfitting to recorder-specific artefacts, background noise, or other spurious signals. By comparing different example mongoose classifications, we can observe the consistency of the model’s focus across similar inputs to detect potential sources of misclassification (see Figure 5).

**Figure 5.**
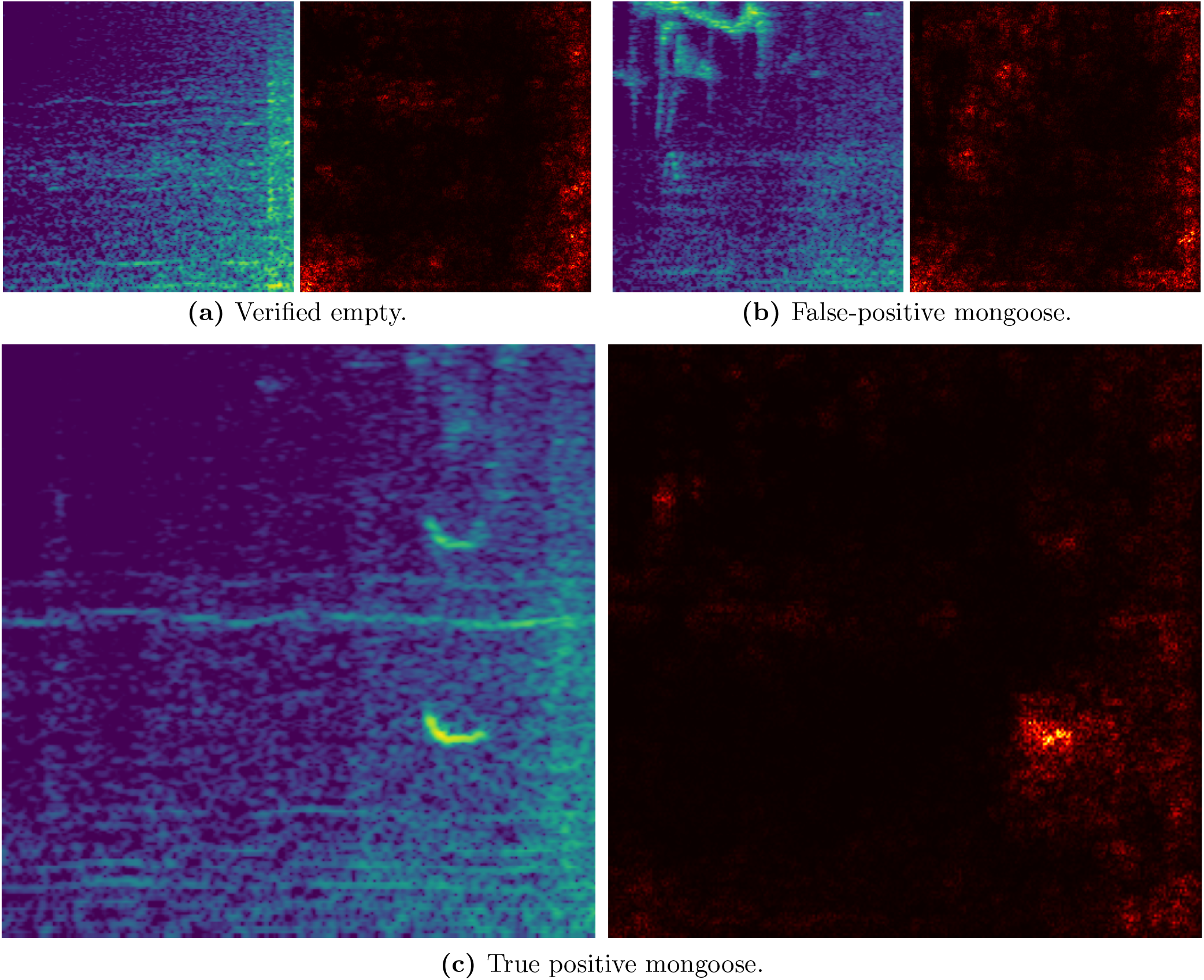
Saliency maps of mongoose classification model. Spectrograms (left) and corresponding saliency maps (right) of predicted (non)mongoose detections. Regions of high saliency (yellow-white) indicate areas that strongly influenced the model’s classification. (a) shows a spectrogram with low confidence (0.119), predicted (and verified) to not contain mongoose. (b) shows a high confidence prediction of mongoose (0.904), but human verification identified it as a brown-eared bulbul *(Hypsipetes amaurotis*); similar frequency regions in the centre left of the activation map may explain the confusion. (c) shows a verified mongoose prediction with high confidence (0.931), with a distinct activated region visible in the saliency map corresponding to the verified mongoose vocalisation. Notably, the harmonic appears largely ignored by the model, suggesting it relies primarily on the base frequency for classification.

## 4 Discussion

Detecting, monitoring, and ultimately controlling the spread of *Urva auropunctata* is of considerable importance for Okinawa’s endemic wildlife threatened by the mongoose’s invasion (Environmental Risk Research Center, National Institute for Environmental Studies, Japan 2025; Yagihashi et al. 2021). We aimed to use audio from camera traps and acoustic recording units in Okinawa as a novel method for detecting and monitoring U. *auropunctata*. This species rarely vocalises, making audio detection a challenge. Indeed, detecting *U. auropunctata* proved challenging, perhaps due to the sparsity of mongoose vocalisations in training and real-world data. To address this, we created a dataset comprising both mongoose vocalisations and negative counter-examples to identify *U. auropunctata* in passive acoustic monitoring data. Using knowledge distillation, we developed a computationally efficient convolutional neural network (CNN) that achieved 92.6% accuracy on the test set. We applied the trained model to a real-world case study—an acoustic dataset spanning eight years—and successfully detected five *U. auropunctata* vocalisations, though this was much fewer than expected. Our approach provides a starting point that has the potential to become a scalable framework for supporting monitoring of this invasive species and hence conservation efforts to protect Okinawa’s unique ecosystems.

Despite the promise of our approach and model performance on test data, we identified several limitations that will need to be overcome before the model can be applied in the field. One key issue is the presence of multiple overlapping vocalisations from other (non-target) animals, which can obscure *U. auropunctata* calls and mislead the model. In particular, the presence of cicada choruses during the summer months poses a challenge for detecting and interpreting acoustic signals due to increased background noise (Ross, Friedman, Yoshimura, et al.2021),and many *U. auropunctata* false positives were detected when both cicada choruses and a non-target species signal were present. Additionally, the model’s performance may be constrained by the limited availability of high-quality, labelled training data. To address these challenges, alternative approaches could be explored, such as building separate classifiers for *U. auropunctata* alarm and contact calls, or countertraining with distinct categories for tricky signals such as the large-billed crow *(Corvus macrorhynchos*, ハシブ ト カラス),brown-eared bulbul *(Hypsipetes amaurotis*,シロガシラ)and Japanese bush warbler *(Horornis diphone*, ウキス). More advanced techniques such as deep metric learning may also be useful to further distinguish different signal classes (Oba and Doi 2025), while simulating soundscapes could expand the limited *U. auropunctata* training set using synthetic samples (Weldy et al. 2025; Gibbons, King, et al. 2024).

Here we discuss several approaches that could enhance model performance in future work:

### (a) Expanded Training Data and Data Augmentation

One of the most reliable ways to improve a classifier’s generalisation is simply to give it more and better examples to learn from. The large Tamagusuku dataset contains validated true positive *U. auropunctata* detections as well as false positives that can serve as hard negative counter-examples. Incorporating these into the training set would increase both data volume and diversity, while providing recorder-specific examples that may help the model adapt to particular hardware characteristics.

Beyond collecting more labelled data, augmentation techniques offer a cost-effective route to a more robust model. Generative augmentation methods (Gibbons, King, et al. 2024; Guei et al. 2024) can synthesise novel call variants, while simulated soundscapes (Weldy et al. 2025) expose the model to realistic combinations of target signals and background noise before it encounters them in the field. Conventional approaches such as SpecAugment (Park et al. 2019), pitch shifting, and mixup, further encourage the model to focus on the core acoustic features of the call rather than incidental recorder artefacts or co-occurring sounds. Together, these strategies directly target the two failure modes most likely to limit field performance: overfitting to a narrow set of recording conditions (Kershenbaum et al. 2025; Lostanlen et al. 2019), and confusion with overlapping environmental signals (Brooker et al. 2020).

### (b) Broadening the Detection Target

A classifier trained only to recognise a single call type is inherently limited by the rarity and variability of that vocalisation. *U. auropunctata* produces both alarm and contact calls, each with distinct temporal and spectral properties; training the model to distinguish these as separate categories, rather than collapsing them into a single positive class, could sharpen decision boundaries and reduce ambiguity when both call types appear in realistic field recordings. A complementary strategy is to include as a separate category of training data alarm calls of prey or disturbed species whose behaviour is reliably altered by mongoose presence (Will Dawson et al. 2026; Gasc et al. 2017). Prey species may vocalise more frequently and conspicuously in response to its predator (i.e., alarm calls) than U. *auropunctata* itself does (Makin, Chamaille-Jammes, and Shrader 2019). The two strategies are not mutually exclusive, and combining a direct call detector with an indirect alarm call detector could yield a more reliable system overall.

### (c) Foundation Model Fine-Tuning

Using model embeddings from BirdNET or Perch (Morfi et al. 2023), rather than the pre-trained models used in this paper, could provide complementary representations of the data for detecting *U. auropunctata*. These are large models specifically trained for bird species classification, whose domain is much closer to mongoose vocalisations than general image or audio clasification models, as we used here. Longer-context models such as NatureLM-audio (Robinson, Sadhu, Vashishth, et al. 2025), which can process extended audio sequences and capture temporal structure across a recording rather than a single short clip, may also be worth exploring. NatureLM-audio has been shown to be effective at zero-shot classification - where the model predicts species it has never been explicitly trained on.

### (d) Self-Supervised Pre-Training

When labelled training data is in short supply, one productive strategy is to first learn general acoustic representations from the much larger pool of *unlabelled* audio from PAM schemes. Selfsupervised methods such as wav2vec 2.0 and AudioMAE achieve this by training a model to solve a pretext (artificial) task — predicting masked portions of a spectrogram, for instance — that requires no human annotation but forces the model to build a rich internal representation of the soundscape before training on the downstream (real) task of interest (Baevski et al. 2020; Huang et al. 2022; Zhao et al. 2024). A model pre-trained in this way on Tamagusuku field recordings would arrive at the fine-tuning stage already familiar with the local acoustic environment. A model starting supervised training already adapted to the general features and artefacts in OKEON audio data should require fewer labelled examples to reach useful performance than training from scratch.

### (e) Agent-Assisted Architecture Search

Selecting an optimal model architecture is itself an optimisation problem, and one that has traditionally required considerable manual experimentation. For example, one might try several CNN depths, convolution kernel and stride sizes, test dropout rates, or experiment with different learning rate schedulers — each requiring a full training run to evaluate. Recent agentic frameworks such as `autoresearch` (Karpathy 2026) automate this process by coupling a large language model (LLM) to a training loop: the model trains for a short fixed interval, its performance on a chosen evaluation metric is recorded, and the agent then proposes architectural modifications before the next training cycle begins. Applied here, this approach would allow the architectures explored in this work to serve as a starting point for a systematic automated search, rather than a fixed endpoint. This optimisation could be directed to prioritise a combination of high precision and recall rather than just aggregate accuracy, given that we wish to find as many of the rarely vocalising *U. auropunctata* as possible.

## 5 Conclusion

Our study demonstrated a field audio-based computational approach for monitoring the small Indian mongoose ( *Urva auropunctata*) in Okinawa. By leveraging sparse vocalisations extracted from camera trap videos collected across the island as part of an ongoing environmental monitoring initiative, we trained a lightweight CNN model to detect *U. auropunctata* sounds. By distilling from a larger model to a smaller model, we achieved comparable (even superior) performance while significantly reducing computational overhead. When applied to ∼486 hours of audio data collected over eight years in Southern Okinawa, the trained classifier could identify U. *auropunctata* vocalisations, but with high false-positive error rates. Our approach has potential for combining sparse audio data with deep learning models to monitor invasive mammals, potentially offering a scalable tool to support conservation efforts aimed at protecting Okinawa’s threatened ecosystems in future. However, additional efforts are first needed to improve model performance, and we outlined some recommendations for future iterations of this model (Figure 6). Applications of a refined classifier to additional locations could further improve our understanding of the spatial distribution of *U. auropunctata*, yielding island-wide insights into mongoose activity, ultimately informing targeted management interventions in Okinawa.

**Figure 6.**
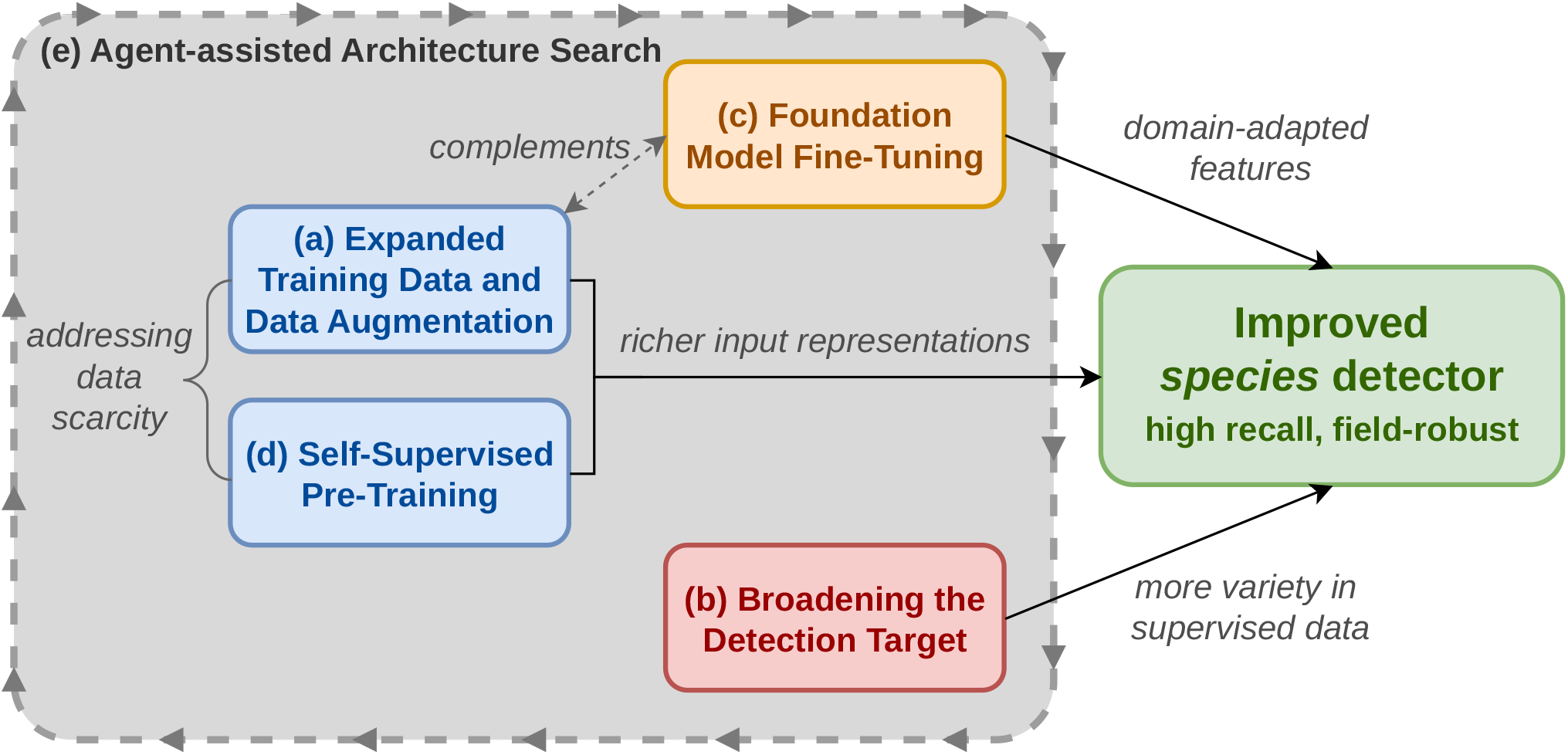
Conceptual overview of proposed improvements to the detection pipeline. Approaches are grouped by their primary point of intervention: data quantity and quality (blue), learned feature representations (orange), and task definition (red). Agent-assisted architecture search (grey) operates as a meta-level optimisation applicable across all model architecture and data input configurations.

## Author Contributions

**A. Gibbons:** Conceptualization, Methodology, Software, Validation, Formal analysis, Investigation, Writing - Original Draft, Data Curation. **I. Donohue:** Writing - Review & Editing, Supervision. **A. Parnell:** Writing - Review & Editing, Supervision, Funding acquisition. **M. Ogasawara:** Resources, Writing - Review & Editing, Funding acquisition. **S.R.P.-J. Ross:** Conceptualization, Methodology, Validation, Investigation, Writing - Original Draft, Visualization, Project administration.

## Acknowledgments

We thank members from the Marine and Terrestrial Science Section of Core Facilities at OIST. We also thank Takumi Akasaka, Sara Keen, and Dave Armitage for early discussion, and Nick Friedman for data included in Figure S1.This work was supported by JST Grant Number JPMJPF2205 and the Nature+Energy Project, funded by Research Ireland (12/RC/2302 P2), industry partners and MaREI, the Research Ireland Research Centre for Energy, Climate and Marine Research and Innovation. MO and SRPJR were supported by subsidy funding to OIST.

## Data and Code Availability

Code and documentation for preprocessing the data, training the species classifiers, inference on large audio dataset, along with the model weights, and post-processing scripts for generating the plots used in this paper are available at https://github.com/gibbona1/OIST_mongoose. All OKEON data, including the raw audio and labelled camera trap data used in this study, are archived with the Okinawa Institute of Science and Technology Graduate University’s high-performance computing center and are available from the Terrestrial and Marine Science Section of Core Facilities at OIST upon request.

## Supplementary Information

### Expanding Dataset via Semi-supervised Learning

We built a feature extractor for semi-supervised learning to find more mongoose vocalisations in the camera trap data. This involved taking the outputs from VGGish and ResNet and training a lightGBM clasifier (Zhang, Si, and Hsieh 2017) to distinguish mongoose from non-mongoose clips. The dataset used consisted of the 32 mongoose clips from 2022 and seven from 2025, and and sample of empty clips from the 2022 dataset (cameras C001 to C025 inclusive). The model was assessed via 5-fold cross validation, achieving ∼80% cross validation accuracy. We then ran the trained model on the remaining 1,600 video clips. We manually checked predictions with > 0.8 confidence above, yielding a further 10 mongoose vocalisations. We then retrained the model with these extra clips and ran the updated model through the entire dataset. We iterated this process and additionally screened data within 5 minutes of validated vocalisations to yield another 10 examples.

### Training and Test Set Creation

The training set was supplemented with examples from the OKEON dataset; these recordings are all from Tamagusuku, the site of the 8-year inference dataset used in this study. This was done for the following reasons:

- **Similar features**. Some examples were misclassified as containing mongoose vocalisations by an earlier version of the model. These examples share certain acoustic features, e.g. bird species vocalising in similar frequency ranges to mongoose, hence the confusion, and so are valuable counter-examples for training the chosen model.
- **Reducing domain shift**. PAM dataset examples should more closely represent the large dataset we run inference on than camera trap audio. In particular, the camera trap audio has more limited spatial coverage than those collecred by the the acoustic recorders.
- **Aligning distribution**. The model, while trained on a roughly 16:1 empty:mongoose ratio, may differ noticeably from the large inference data. The inference data may have much less than 1% that contain mongoose vocalisations. Adding high-quality negative samples should align more features with empty clips, reducing false positives.

Once the training data was constructed, examples were set to be 3 seconds from the annotation start, which biases towards the mongoose vocalisation being at the beginning of the examples. Clips

