## Supplementary material for "Acoustic detection of a rarely vocalising invasive mammal from sparse data": _

Zhang, Tao et al. (2021). "Acoustic scene classification based on Mel spectrogram decomposition and model merging". In: *Applied Acoustics* 182, p. 108258. ISSN: 0003-682X. DOI: https://doi.org/10.1016/j.apacoust.2021.108258.

Zhao, Zehui et al. (2024). "A comparison review of transfer learning and self-supervised learning: Definitions, applications, advantages and limitations". In: *Expert Systems with Applications* 242, p. 122807. ISSN: 0957-4174. DOI: 10.1016/j.eswa.2023.122807.

Zhong, Ming et al. (2020). "Multispecies bioacoustic classification using transfer learning of deep convolutional neural networks with pseudo-labeling". In: *Applied Acoustics* 166, p. 107375. ISSN: 0003-682X. DOI: 10.1016/j.apacoust.2020.107375.

Zulko, Rémy et al. (2025). *MoviePy: Video editing with Python*. Version 2.0. URL: https://github.com/Zulko/moviepy.

### Supplementary Information

#### Expanding Dataset via Semi-supervised Learning

We built a feature extractor for semi-supervised learning to find more mongoose vocalisations in the camera trap data. This involved taking the outputs from VGGish and ResNet and training a lightGBM clasifier (Zhang, Si, and Hsieh 2017) to distinguish mongoose from non-mongoose clips. The dataset used consisted of the 32 mongoose clips from 2022 and seven from 2025, and and sample of empty clips from the 2022 dataset (cameras C001 to C025 inclusive). The model was assessed via 5-fold cross validation, achieving ∼80% cross validation accuracy. We then ran the trained model on the remaining 1,600 video clips. We manually checked predictions with $> 0.8$ confidence above, yielding a further 10 mongoose vocalisations. We then retrained the model with these extra clips and ran the updated model through the entire dataset. We iterated this process and additionally screened data within 5 minutes of validated vocalisations to yield another 10 examples.

#### Training and Test Set Creation

The training set was supplemented with examples from the OKEON dataset; these recordings are all from Tamagusuku, the site of the 8-year inference dataset used in this study. This was done for the following reasons:

- **Similar features.** Some examples were misclassified as containing mongoose vocalisations by an earlier version of the model. These examples share certain acoustic features, e.g. bird species vocalising in similar frequency ranges to mongoose, hence the confusion, and so are valuable counter-examples for training the chosen model.
- **Reducing domain shift.** PAM dataset examples should more closely represent the large dataset we run inference on than camera trap audio. In particular, the camera trap audio has more limited spatial coverage than those collecred by the the acoustic recorders.
- **Aligning distribution.** The model, while trained on a roughly 16:1 empty:mongoose ratio, may differ noticeably from the large inference data. The inference data may have much less than 1% that contain mongoose vocalisations. Adding high-quality negative samples should align more features with empty clips, reducing false positives.

Once the training data was constructed, examples were set to be 3 seconds from the annotation start, which biases towards the mongoose vocalisation being at the beginning of the examples. Clips were randomly ‘rolled’ so the vocalisation(s) present occur at different points in the clip. Average waveforms were analysed for stereo audio files. The signal was converted to a mel-spectrogram with `n_fft`=4096, `hop_length`=512, `n_mels`=256, `power`=4.0 and cropped to 400 Hz–9 kHz. The output spectrograms were cropped to 256×256 single-channel (grayscale) and repeated to three channels for input to ResNet. To reduce background noise, each spectrogram had the mean of each frequency (y-axis) bin subtracted across the entire row. Finally, the spectrograms were normalised to be in the range [0,1] and then normalised again with the means and standard deviations from ImageNet dataset, to conform with the input format the pre-trained ResNet expects. VGGish uses the 3-second signal array and its sample rate as inputs.

#### Classification Model Construction

The initial model for mongoose classification made use of two pre-trained CNNs. The first is VGGish (Hershey et al. 2017), a VGG-like architecture trained on AudioSet (Gemmeke et al. 2017). It takes raw 3-second mono audio and its sample rate as input and generates feature embeddings for each second, which are then flattened to a single vector of length $128 \times 3 = 384$. Concurrently, a ResNet model (He et al. 2015), pre-trained on ImageNet (Deng et al. 2009), takes in a 256×256×3 image and outputs an embedding of length 1000. All but the final layer of the ResNet model were frozen; the model can update the unfrozen layer during training. The embeddings from both model outputs are concatenated to a vector of length 1384 and passed through 3 fully connected (FC) layers, with dropout with probability 0.5 applied after each to reduce overfitting. Using two feature extractors trained on different datasets and modalities (images for ResNet, audio for VGGish) provides complementary insights from both models. During training, the VGGish embeddings and mel spectrograms can be saved after their initial generation in the first epoch and loaded in future epochs, reducing training time. The ResNet embedding changes since the final layer is trainable, but an earlier layer could technically have been saved.

Our target dataset consists of over ∼486 hours of data, requiring ∼583,000 spectrograms (and waveform arrays for VGGish) to be computed and run through the model. A smaller CNN, while faster than the above, may not be able to generalise given the relatively small size of the mongoose audio dataset. To address these computational constraints, while also taking advantage of the learnings of the two pre-trained models, the second iteration of the classifier is a smaller CNN which takes in only spectrograms and does not make direct use of the pre-trained classifiers. Using knowledge distillation, where the larger model from the first iteration serves as the teacher (Hinton, Vinyals, and Dean 2015) for the smaller student model, the training objective combines cross-entropy loss, as in the teacher model, with Kullback-Leibler (KL) divergence-based distillation loss to align the student model’s softmax outputs with the teacher’s. Since the output has only two classes, this may not be enough to transfer much of the learnings from teacher to student. The distilled student model was trained additionally with a cosine embedding loss aligning outputs from an earlier layer (with dimension 256) in both models. This enables the smaller model to retain much of the teacher’s predictive power while reducing parameters and inference time. A comparison of the teacher and smaller student model used in this paper is given in Table 1.

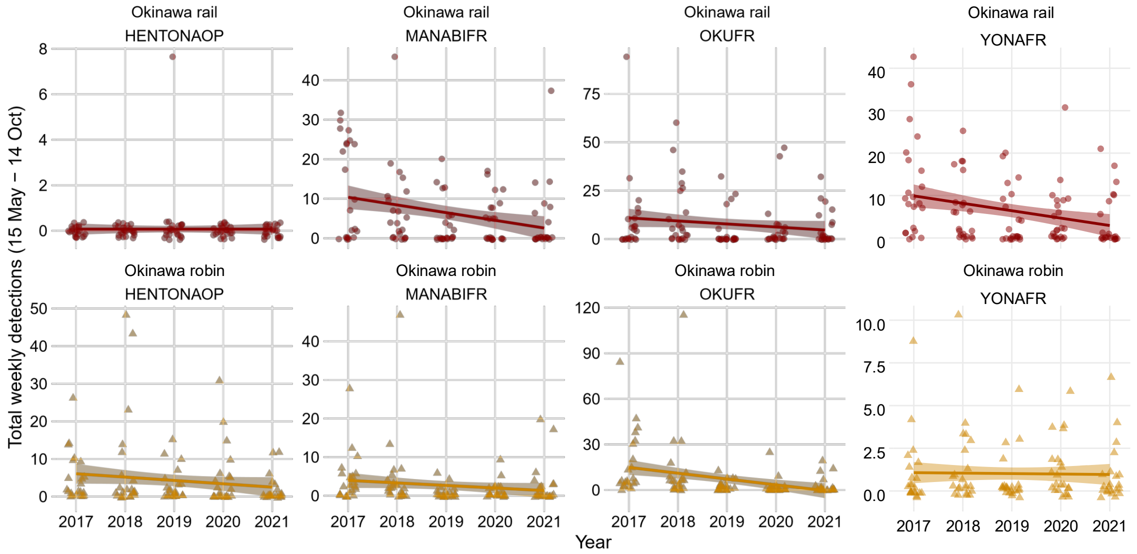


**Figure S1: Detection trends for Yambaru bird species.** Total weekly detections for the 15 May-14 October period per year spanning 2017-2021. Data is derived from a BirdNET model (unpublished data), but shows declines in both the Okinawa rail and Okinawa robin in most cases.

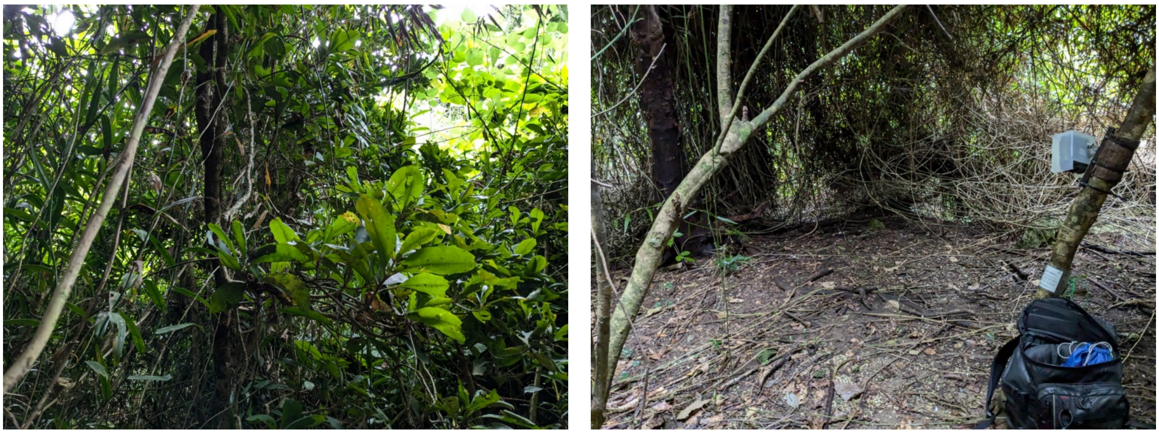

**(a)** Tamagusuku canopy.

**(b)** Tamagusuku forest floor.

**Figure S2: Tamagusuku site.** Photo credit: Damien Morrell

#### Figures

#### Web-scraped data

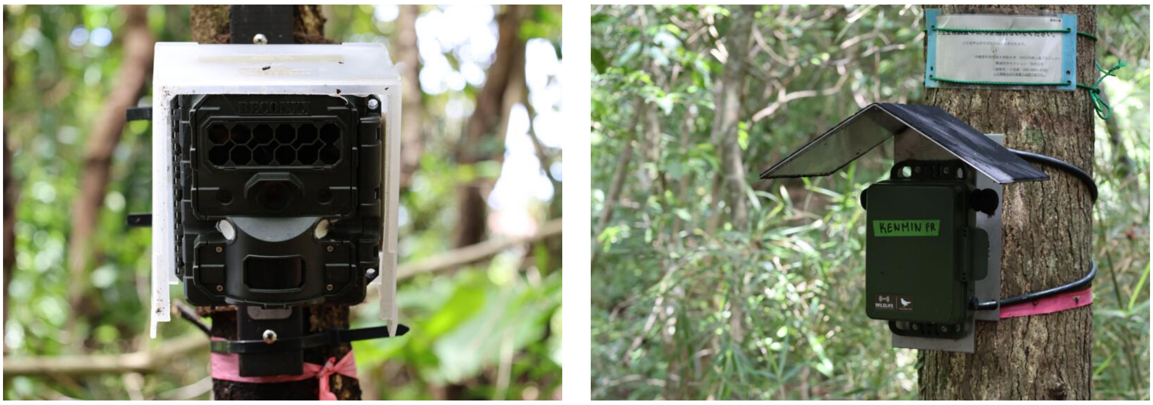

**(a)** Camera Trap. **(b)** Acoustic Recorder.

**Figure S3: OKEON recording equipment.**

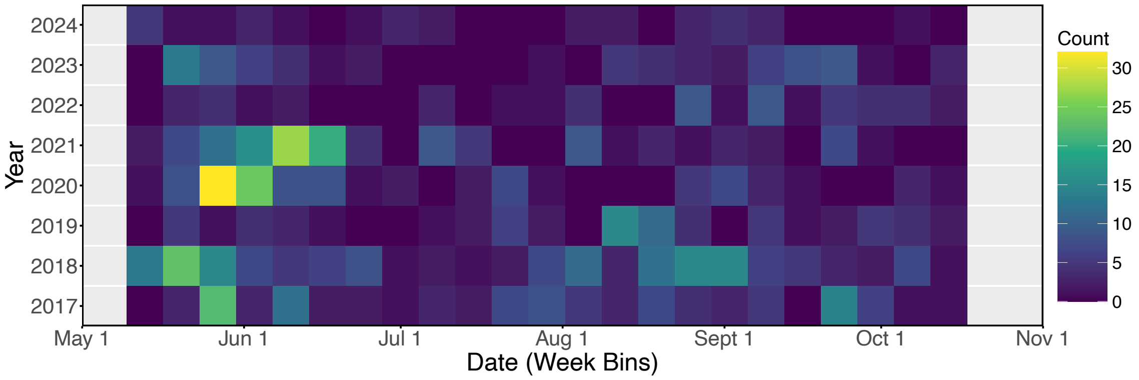


**Figure S4: Counts of confident model predictions over the dataset length.**


Gemmeke, Jort F. et al. (2017). "Audio Set: An ontology and human-labeled dataset for audio events". In: *2017 IEEE International Conference on Acoustics, Speech and Signal Processing (ICASSP)*, pp. 776–780. DOI: 10.1109/ICASSP.2017.7952261.

He, Kaiming et al. (2015). *Deep Residual Learning for Image Recognition.* URL: https://arxiv.org/abs/1512.03385.

Hershey, Shawn et al. (2017). *CNN Architectures for Large-Scale Audio Classification.* URL: https://arxiv.org/abs/1609.09430.

Hinton, Geoffrey E, Oriol Vinyals, and Jeff Dean (2015). *Distilling the Knowledge in a Neural Network.* URL: https://arxiv.org/abs/1503.02531.

Zhang, Huan, Si Si, and Cho-Jui Hsieh (2017). *GPU-acceleration for Large-scale Tree Boosting.* URL: https://arxiv.org/abs/1706.08359.

| Filename | URL | Length (s) |
|---|---|---|
| short1.wav | https://youtu.be/AE9BEdvXfF8 | 18 |
| short2.wav | https://youtu.be/rtN1NoXM3Cc | 7 |
| short3.wav | https://youtu.be/-mOD-sDuWS4 | 10 |
| video1.wav | https://youtu.be/V2QP1bQSSc0 | 9 |
| video2.wav | https://youtu.be/jdvEn9VxyQg | 15 |
| video3.wav | https://youtu.be/ch5-exHYlG4 | 34 |
| video4.wav | https://youtu.be/2smtcRvugnU | 82 |
| video5.wav | https://youtu.be/ISWCrjGZVjw | 67 |
| video6.wav | https://facebook.com/share/v/1BN4bVzecR | 15 |
| XC578808.wav | https://xeno-canto.org/578808 | 153 |
| XC156050.wav | https://xeno-canto.org/156050 | 109 |
| XC156097.wav | https://xeno-canto.org/156097 | 71 |
| XC156111.wav | https://xeno-canto.org/156111 | 23 |
| XC191039.wav | https://xeno-canto.org/191039 | 66 |
| XC202788.wav | https://xeno-canto.org/202788 | 6 |
| XC202790.wav | https://xeno-canto.org/202790 | 16 |
| XC236034.wav | https://xeno-canto.org/236034 | 20 |
| XC286152.wav | https://xeno-canto.org/286152 | 17 |
| XC474444.wav | https://xeno-canto.org/474444 | 59 |
| XC578817.wav | https://xeno-canto.org/578817 | 3 |
| XC884889.wav | https://xeno-canto.org/884889 | 56 |
| XC984255.wav | https://xeno-canto.org/984255 | 42 |
| XC984256.wav | https://xeno-canto.org/984256 | 43 |

**Table S1: Web scraped audio details.** The web scraped audio was extracted from Xeno-Canto recordings and YouTube and Facebook videos. The two Xeno-Canto files beginning '98' are of the Crab-eating Mongoose recorded in Taiwan, but the vocalisations were deemed to be similar enough to the species of interest, *Urva auropunctata*. The remaining Xeno-Canto files were bird species used as counter-examples for training. The majority of the mongoose recordings are alarm calls, while short1 and video6 contain contact calls. In the final training set, these web-scraped examples constitute nearly half of the mongoose examples (these were excluded from the validation and test sets), underscoring their utility in enhancing the dataset's diversity and improving model robustness.
